# The BRPF1 bromodomain is a molecular reader of di-acetyllysine

**DOI:** 10.1101/2020.02.13.948091

**Authors:** Juliet O. Obi, Mulu Y. Lubula, Gabriel Cornilescu, Amy Henrickson, Kara McGuire, Chiara M. Evans, Margaret Phillips, Samuel P. Boyson, Borries Demeler, John L. Markley, Karen C. Glass

## Abstract

Bromodomain-containing proteins are often part of chromatin-modifying complexes, and their activity can lead to altered expression of genes that drive cancer, inflammation and neurological disorders in humans. Bromodomain-PHD finger protein 1 (BRPF1) is part of the MOZ (monocytic leukemic zinc-finger protein) HAT (histone acetyltransferase) complex, which is associated with chromosomal translocations known to contribute to the development of acute myeloid leukemia (AML). BRPF1 contains a unique combination of chromatin reader domains including two plant homeodomain (PHD) fingers separated by a zinc knuckle (PZP domain), a bromodomain, and a proline-tryptophan-tryptophan-proline (PWWP) domain. BRPF1 is known to recruit the MOZ HAT complex to chromatin by recognizing acetylated lysine residues on the N-terminal histone tail region through its bromodomain. However, histone proteins can contain several acetylation modifications on their N-terminus, and it is unknown how additional marks influence bromodomain recruitment to chromatin. Here, we identify the BRPF1 bromodomain as a selective reader of di-acetyllysine modifications on histone H4. We used ITC assays to characterize the binding of di-acetylated histone ligands to the BRPF1 bromodomain and found that the domain binds preferentially to histone peptides H4K5acK8ac and H4K5acK12ac. Analytical ultracentrifugation (AUC) experiments revealed that the monomeric state of the BRPF1 bromodomain coordinates di-acetylated histone ligands. NMR chemical shift perturbation studies, along with binding and mutational analyses, revealed non-canonical regions of the bromodomain-binding pocket that are important for histone tail recognition. Together, our findings provide critical information on how the combinatorial action of post-translational modifications can modulate BRPF1 bromodomain binding and specificity.

## INTRODUCTION

In the human genome, DNA accessibility and transcriptional activation are often regulated by post-translational modifications (PTMs), among which acetylation of lysine residues on histones plays a major role. Histone acetylation enhances DNA accessibility by loosening the compact conformation of chromatin and by recruiting transcription and chromatin remodeling factors to increase transcriptional activity^1,2^. Bromodomains, which are epigenetic reader domains that specifically recognize ε-N-acetylated lysine residues on histone tails, are known to be important regulators of chromatin remodeling and transcriptional activation^3^. A total of 61 unique bromodomains are known to exist in 46 different bromodomain-containing proteins, and these bromodomains have been grouped into eight major families on the basis of sequence and structural homology^4^. Mutation and aberrant expression of bromodomain-containing proteins has been associated with multiple types of cancer, inflammation, neurological disorders and metabolic diseases. Importantly, bromodomains are often found in chromatin-modifying complexes, whose activity can lead to the altered expression of genes that drive these diseases in humans^5^.

Bromodomain-PHD finger protein 1 (BRPF1) is known to play a role in maintaining the expression of genes involved in several developmental processes and hematopoiesis^6-8^. The BPRF1 protein is a unique, multivalent chromatin reader that contains an N-terminal plant homeodomain (PHD)-zinc-knuckle-PHD (PZP) module, a bromodomain for acetyllysine recognition, and a C-terminal proline-tryptophan-tryptophan-proline (PWWP) domain^9^. The N-terminal motif I of BRPF1 interacts with monocytic leukemic zinc finger (MOZ) and MOZ-related factor (MORF), while the downstream motif II interacts with inhibitor of growth 5 (ING5) and the MYST/Esa1-associated factor 6 (MEAF6)^10^. Thus, an important function of BRPF1 is to bridge the association of MOZ and MORF with ING5 and MEAF6, where it is an important regulator of many cellular processes including the self-renewal of leukemic stem cells^8^, embryo development and cell proliferation^11^, and osteoclast differentiation^12^ through its activity with MOZ and MORF^8^. BRPF1 enhances the acetyltransferase activity of the MOZ/MORF HATs via interaction with the histone acetyltransferase domain^10^, and also bridges the enzymatic complex that carries out specific post-translational modifications in chromatin^13^.

Post-translational modifications (PTMs) on histones are known to be crucial for the regulation of chromatin and DNA accessibility^14^. PTMs modulate signal transduction pathways to recruit effector molecules, such as readers of histone modifications to chromatin^15^. These histone reader domains direct biological outcomes based on the cellular context of the modifications^16^. In humans, proteins are known to undergo a number of PTMs, including but not limited to phosphorylation, methylation, acetylation and ubiquitination, that greatly influence their function in cellular events^17^. Specifically, histone acetylation is an important component of epigenetic signaling, in that it weakens the interactions of histones with DNA by decreasing nucleosome compaction^18^, regulates enzymatic activity, and mediates the localization of macromolecular complexes to histones through bromodomain-containing proteins^19^. Therefore, the recognition of acetylated lysine by bromodomains has broad implications for cellular events including chromatin remodeling and transcriptional activation^20^.

The postulation that aberrant chromatin acetylation by MOZ may be a contributing factor to the development of cancer and other diseases was first demonstrated by cloning the breakpoint-associated genes of the recurrent t(8;16)(p11;p13) translocation in patients with a subtype of acute myeloid leukemia (AML)^21^. This translocation, which results in a novel fusion that combines the PHD finger motifs and MYST acetyltransferase domain of MOZ with an intact CREB-binding protein (CBP), suggested that the MOZ-CBP fusion protein gives rise to leukemogenesis through chromatin hyperacetylation^21^. Since BRPF1 plays a crucial role in the MOZ/MORF complex, and the interaction domains of this complex remain intact after the translocation, interaction with BRPF1 may be implicated in other diseases in which MOZ and MORF are known to play a role. The *MOZ* gene has been reported to be mutated in esophageal adenocarcinoma^22^, while the *MORF* gene has been reported to be disrupted in leiomyomata^23,24^, mutated in familial breast cancer^25^, and altered in castration-resistant prostate cancer^26^. Deletion of the mouse *BRPF1* gene showed that BRPF1 is crucial for the growth and proliferation of embryonic fibroblasts and hematopoietic progenitors^11^. Supporting evidence that BRPF1 acts through MOZ and MORF to modulate chromatin modification states comes from the report that multiple mutations in the *BRPF1* gene have been reported to give rise to neurodevelopmental disorders associated with a deficiency in histone H3K23 acetylation^7^.

The BRPF1 bromodomain belongs to the family IV bromodomains, which also include the BRPF2, BRPF3, BRD7, BRD9, ATAD2 and ATAD2B bromodomains based on their sequence and structural homology^4^. Bromodomains share a conserved globular fold and are composed of a left-handed bundle of four α-helices Z, A, B and C. The helices are linked by loops of variable length (ZA and BC loops), which form the hydrophobic acetyllysine (Kac) binding pocket^4^. Differences in teh acetyllysine recognition of these bromodomains is attributed mainly due to the variability of their ZA and BC loops^4^. These differences, have also permitted the development of small molecule inhibitors with selectivity for specific bromodomains within the same family^27-30^.

An earlier study characterized the interaction of the BRPF1 bromodomain with mono-acetylated ligands on the N-terminal of histone tails, and showed selectivity for multiple acetylation modifications including H2AK5ac, H4K12ac and H3K14ac^31^. However, each of these four histone proteins can be post-translationally modified at multiple locations, and with several different chemical marks^32^. Proteomic studies have demonstrated that each histone tail caries a unique set of modifications^33^, and that the abundance and availability of specific histone proteoforms is dynamic^34,35^. The BET bromodomains BRD2, BRD4 and BRDT are all able to bind di-acetyllysine modifications including H4K5acK8ac, H4K5acK12ac, and H4K12acK16ac^4,36^. Other family IV bromodomains, including the BRD9 and ATAD2 bromodomain, are able to interact with di-acetylated histone ligands such as H4K5acK8ac and H4K5acK12ac, respectively^19,37^. However, the interaction of the BRPF1 bromodomain with di-acetyllysine modifications has not been systematically characterized, and it is unknown how multiple histone marks modulate acetyllysine recognition.

Previously published data on the BRPF1 bromodomain revealed that it recognizes several acetylated lysine residues on N-terminal histone tails^31^. Related bromodomain proteins have been observed to cooperatively bind histone peptides with di-acetyllysine marks, through accommodation of the two acetyllysine residues in the same bromodomain binding pocket. For example, the BRDT bromodomain BD1 in the bromodomain and extra-terminal domain (BET) family, recognizes the di-acetylated histone ligand H4K5acK8ac^36^. Other bromodomain proteins in the BET family including BRD2 and BRD4 are also known to recognize di-acetylated histone ligands, mainly through their enlarged binding pockets^4^. Three-dimensional X-ray crystal structures of the BAZ2A and BAZ2B bromodomains in family V have also been solved in complex with H4K16acK20ac and H4K8acK12ac di-acetylated histone ligands^38^. The family IV bromodomains possess a “keyhole style” pocket due to a larger aromatic gatekeeper residue that can only accommodate one acetyllysine group in the canonical binding site^19,36^. In ITC binding assays and crystallization studies of BRD9, the bromodomain interacts with di-acetylated histone ligand by coordinating the H4K5ac group in one bromodomain with the H4K8ac site bound to a second bromodomain module (PDBID: 4YYI)^19^. However, the binding and recognition of di-acetylated histone ligands by the BRPF1 bromodomain has yet to be fully elucidated.

In this study we characterized the binding affinity of the BRPF1 bromodomain with multiple di-acetylated histone ligands by isothermal titration calorimetry (ITC). We show that the BRPF1 bromodomain binds preferentially to histone peptides H4K5acK8ac and H4K5acK12ac. We utilized nuclear magnetic resonance (NMR) chemical shift perturbation techniques to map the BRPF1 bromodomain binding pocket and identify important residues involved in di-acetyllysine recognition. We performed site-directed mutagenesis experiments, together with ITC and analytical ultracentrifugation (AUC) experiments to evaluate the relative contributions of specific amino acid residues in di-acetyllysine coordination. We also tested how adjacent PTMs impact acetylysine recognition. The results provide new insights on how the BRPF1 bromodomain selects for its di-acetyllysine histone ligands at the molecular level, and how the combinatorial action of post-translational modifications can influence BRPF1 bromodomain binding and specificity. Our study further supports current research efforts geared toward the development of novel therapeutics targeting the BRPF1 bromodomain in acute myeloid leukemia.

## RESULTS

### The BRPF1 bromodomain recognizes di-acetyllysine histone modifications

We postulated that the BRPF1 bromodomain may be able to recognize di-acetylated histone ligands in analogy to the closely related family IV bromodomain BRD9, which has been shown to recognize multiple diacyl modifications on histone H4K5/K8^19^. To investigate our hypothesis, we sought to further characterize the histone ligands of this bromodomain by performing ITC experiments with mono- and di-acetylated lysine residues on the histones H3 and H4. The BRPF1 bromodomain is able to recognize several acetyllysine modifications, and binds to several mono-acetylated histone peptides, showing highest affinity toward the H4K5ac peptides (residues 1-10 and 1-15) (**Table 1 and Supplementary Figure S1)**. Furthermore, when the ITC experiment was performed with the di-acetylated H4K5acK8ac (1-10) peptide, the BRPF1 bromodomain bound with a dissociation constant about two-fold stronger than that of the mono-acetylated H4K5ac peptide (27.7 ± 5.2 µM versus 57.6 ± 6.0). Similarly, we found that the combination of H4K5acK12ac marks increased the BRPF1 bromodomain binding affinity to a K_D_ of 49.7 ± 6.5 µM, six-fold over that for the singly acetylated H4K12ac mark, and has a higher affinity than the single H4K5ac modification. Other combinations of di-acetylation marks on the histone tail resulted in lower binding affinities with the BRPF1 bromodomain, and the H4K8acK12ac mark completely abrogated the binding interaction.

**Table 1:** Sequences of the mono- and di-acetylated histone peptides studied, and the dissociation constants of their complexes with the BRPF1 bromodomain as measured by ITC.

| Histone peptide | Sequence | BRPF1<br>K<sub>D</sub> (μM) | N value | ΔH<br>(cal/Mol) | ΔS<br>(cal/Mol/deg) |
| --- | --- | --- | --- | --- | --- |
| **Histone H3 peptides** |  |  |  |  |  |
| H3 unmodified (1–24) | ARTKQTARKSTGGK<br>APRKQLATKA | No binding | --- | --- | --- |
| H3K9ac (1–24) | ARTKQTAR(Kac)STG<br>GKAPRKQLATKA | 718.0 ±<br>55.8 | 1.0 ± 0.0 | -918.8 ± 151.13 | 11.1 ± 0.4 |
| H3K14ac (1–24) | ARTKQTARKSTGG(<br>Kac)APRKQLATKA | 183.7 ± 3.9 | 1.0 ± 0.0 | -1338.5 ± 48.7 | 12.3 ± 0.2 |
| H3K14ac (8–19)<sup>1</sup> | RKSTGG(Kac)APRKQ | 626 | --- | --- | --- |
| H3K9acK14ac (1–24) | ARTKQTAR(Kac)STG<br>G(Kac)APRKQLATK<br>A | 233.8 ± 5.8 | 1.0 ± 0.0 | -1249.3 ± 136.7 | 12.1 ± 0.5 |
| **Histone H4 peptides** |  |  |  |  |  |
| **Mono-acetylated histone H4 peptides** |  |  |  |  |  |
| H4 unmodified (1–15) | SGRGKGGKGLGKG<br>GA | No binding | --- | --- | --- |
| H4K5ac (1–10) | SGRG(Kac)GGKGL | 57.6 ± 6.0 | 0.9 ± 0.1 | -3823.0 ± 220.0 | 5.7 ± 0.7 |
| H4K5ac (1–15) | SGRG(Kac)GGKGLG<br>KGGA | 78.7 ± 15.4 | 1.0 ± 0.1 | -3487.3 ± 450.1 | 6.3 ± 1.3 |
| H4K5ac (4–17)<sup>1</sup> | G(Kac)GGKGLGKGG<br>AKR | 1236* | --- | --- | --- |
| H4K8ac (1–10) | SGRGKGG(Kac)GL | 189.1 ±<br>36.6 | 1.1 ± 0.2 | -622.5 ± 110.1 | 14.8 ± 0.1 |
| H4K8ac (4–17)<sup>1</sup> | GKGG(Kac)GLGKGG<br>AKR | 697* | --- | --- | --- |
| H4K12ac (1–15) | SGRGKGGKGLG(Kac<br>)GGA | 100.0 ± 4.6 | 1.0 ± 0.1 | -2544.7 ± 430.0 | 9.0 ± 1.3 |
| H4K12ac (4–17)<sup>1</sup> | GKGGKGLG(Kac)GG<br>AKR | 86.5 ± 9.1 | --- | --- | --- |
| H4K16ac (10–20) | LGKGGA(Kac)RHRK | 517.9 ±<br>92.1 | 1.1 ± 0.1 | -3862.7 ± 112.0 | 1.2 ± 0.1 |
| **Di-acetylated histone H4 peptides** |  |  |  |  |  |
| H4K5acK8ac (1–10) | SGRG(Kac)GG(Kac)G<br>L | 27.7 ± 5.2 | 0.8 ± 0.0 | -1186 ± 35.3 | 16.6 ± 0.3 |
| H4K5acK12ac (1–15) | SGRG(Kac)GGKGLG(<br>Kac)GGA | 49.7 ± 6.5 | 1.0 ± 0.0 | -7216.0 ± 378.5 | -6.2 ± 1.2 |
| H4K5acK16ac (1–24) | SGRG(Kac)GGKGLG<br>KGGA(Kac)RHRKVL<br>RD | 981 ± 11.3 | 1.0 ± 0.1 | -4560.0 ± 148.8 | 2.0 ± 0.3 |
| H4K8acK12ac (6–15) | GG(Kac)GLG(Kac)GG<br>A | No binding | --- | --- | --- |
| H4K8acK16ac (1–24) | SGRGKGG(Kac)GLG<br>KGGA(Kac)RHRKVL<br>RD | 444.7 ±<br>12.5 | 1.0 ± 0.0 | -3844.3 ± 56.0 | 1.5 ± 0.2 |
| H4K12acK16ac (10–<br>20) | LG(Kac)GGA(Kac)RH<br>RK | 644.2 ±<br>102.3 | 1.0 ± 0.0 | -4577.7 ± 179.4 | 1.8 ± 0.3 |
| **Other PTMs on histone H4 peptides** |  |  |  |  |  |
| H4S1phosK5ac (1–15) | (Sp)GRG(Kac)GGKGL<br>GKGGA | 220.3 ±<br>25.2 | 1.1 ± 0.1 | -3523.3 ± 195.3 | 4.1 ± 0.5 |
| H4K5me3K12ac (1–15) | SGRG(Kme3)GGKGL<br>G(Kac)GGA | 179.8 ±<br>13.4 | 1.0 ± 0.0 | -4294.0 ± 64.3 | 1.7 ± 0.1 |
| H4K5acK12me3 (1–15) | SGRG(Kac)GGKGLG(<br>Kme3)GGA | 70.3 ± 4.8 | 1.1 ± 0.0 | -6357.0 ± 205.9 | -3.8 ± 0.6 |
| <sup>1</sup> Previously published BRPF1 binding data<sup>31</sup> |  |  |  |  |  |
| * Indicates dissociation constants measured by NMR titration experiments |  |  |  |  |  |

Interestingly, in some cases, inclusion of residues 1-3 on the histone tail improved the binding interaction of histone H4 peptides that contained the K5 or K8 acetyllysine modification. This is the case with H4K5ac peptides (comparison of residues 1-10 and 4-17), and H4K8ac peptides (comparison of residues 1-10 and 4-17), bind with significantly diffferent affinities (**Table 1**)^31^. However, in peptides with the H4K12ac modification, sequences 1-15 and 4-17 had similar binding constants. These data indicate that residues 1-3 on the N-terminus of histone H4 tail are important for recognition of acetyllysine modifications near the N-terminus of this histone, whereas the presence of N-terminal residues has little effect on the affinity of modifications further downstream. Furthermore, this is the first direct evidence that the BRPF1 bromodomain can read a span of 8-10 amino acid residues on the histone tail.

Since previous studies have shown that the BRPF1 bromodomain recognizes the H3K14ac histone peptide^31^, we also tested the binding affinity of this bromodomain with mono- and di-acetylated peptides containing the first 24 amino acid residues of human histone H3. Our ITC experiments revealed that the BRPF1 bromodomian preferentially selects for the H3K14ac (1-24) modification over H3K9ac (1-24). A dissociation constant of 233.8 µM was obtained for the H3K9acK14ac (1-24) di-acetylated histone peptide, which indicates that the acetylation mark at K9 impairs the ability of the BRPF1 bromodomain to bind H3K14ac (**Table 1, Supplementary Figure S1).**

We calculated ΔH and ΔS values for the interaction of BRPF1 with various mono- and di-acetylated histone ligands (**Table 1).** The bromodomain-histone binding interaction appears to be predominantly enthalpy driven as demonstrated by the large, negative changes in enthalpy (ΔH) observed upon ligand binding. This is opposition to the smaller, generally positive changes in entropy (ΔS), which may reflect the displacement of water molecules in the bromodomain binding pocket upon ligand binding^39^. Taken together, our results show that the BRPF1 bromodomain preferentially selects for di-acetyllysine modifications on histone H4.

### Crosstalk between PTMs modulate acetyllysine recognition by the BRPF1 bromodomain

To determine how the combinatorial action of PTMs on the histone tail modulates acetyllysine recognition, we performed ITC experiments to measure how the *in vitro* binding affinity of the BRPF1 bromodomain for acetylated histone peptides changes as a function of modifications located adjacent to the acetyllysine (Kac) recognition site (**Table 1 and Supplementary Figure S1**). Our data show that recognition of the H4K5ac (1-15) mono-acetylated histone peptide decreased by approximately three-fold (K_D_ = 228.5 µM) when serine 1 of histone H4 was phosphorylated. Additionally, the affinity of the H4K12ac (1-15) interaction decreased by ∼1.5-fold (169.6 µM) when lysine 5 was tri-methylated. Interestingly, tri-methylation of lysine 12 in combination with acetylation of lysine 5 did not significantly alter the affinity of the BRPF1 bromodomain for H4K5ac (1-15), indicating that the H4K12me3 mark is permissive for H4K5ac recognition. Together, our results show that crosstalk between adjacent PTMs on the histone tail modulate how acetyllysine is recognized by the BRPF1 bromodomain.

### The BRPF1 bromodomain recognizes di-acetylated histone ligands as a monomer

In a recent crystal structure, the complex formed between the BRPF1 bromodomain and histone H4K5acK8ac is modeled as an apparent dimer (PDBID: 5FFW)^12^. However, we hypothesized that BRPF1 likely functions as a monomer in its biologically active form when it associates with the MOZ/MORF HATs. As shown in **Table 1**, the N value observed in all our ITC data indicates that the BRPF1 bromodomain interacts with modified histone H3 and H4 tails as a monomer. To confirm this, we also performed analytical ultracentrifugation sedimentation velocity experiments to determine the molecular masses of the BRPF1 bromodomain in complexes with mono- and di-acetylated histone ligands. The molecular weight of the apo BRPF1 bromodomain is 13,703 Da, while the molecular weights of the histone ligands tested ranged from 958 to 1371 Da. Strikingly, the signal peaks of the BRPF1 bromodomain alone, or in complex with its mono- and di-acetylated histone ligands, overlaped at approximately 14-15 kDa (**Figure 1**). These results support our ITC data showing that the BRPF1 bromodomain recognizes both mono- and di-acetylated histone ligands as a monomer in solution.

**Figure 1.**
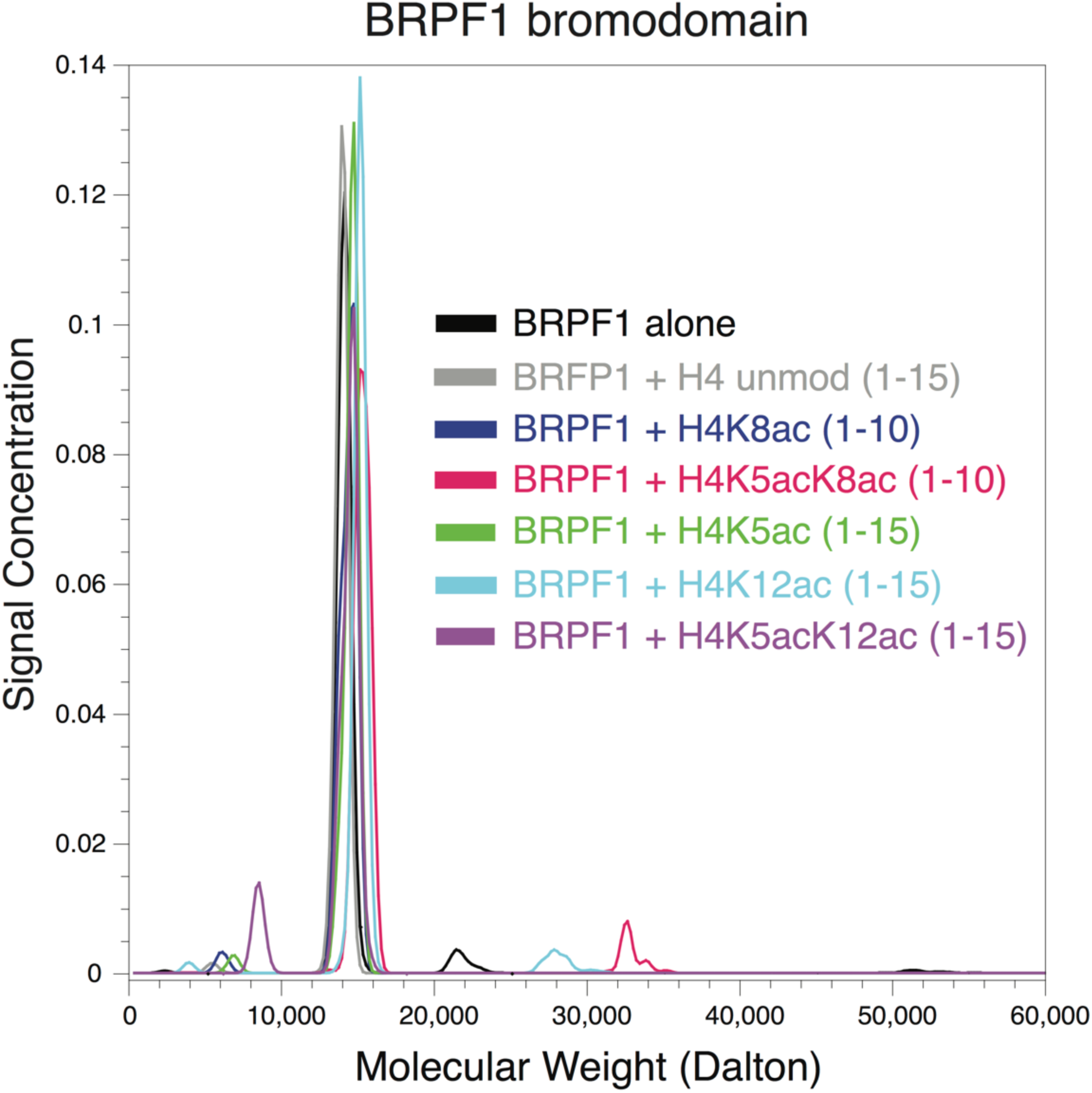
Molecular weight distributions of the BRPF1 bromodomain alone and in complex with mono- and di-acetylated histone ligands, determined by analytical ultracentrifugation. The distribution shows the molecular weights of the BRPF1 bromodomain alone and a 1:1 combination ratio with H4K5ac (1-15), H4K5acK12ac (1-15), H4K12ac (1-15), H4 unmodified (1-15), H4K5acK8ac (1-10), and H4K8ac (1-10) colored in red, blue, magenta, purple, green, black, and cyan respectively. All molar masses are consistent with a 1:1 ratio of BRPF1 bromodomain and histone peptide.

### Characterization of the BRPF1 bromodomain binding pocket by NMR spectroscopy

To outline specific amino acid interactions in the BRPF1 bromodomain binding pocket with the mono- and di-acetylated histone peptides, two-dimensional, ^1^H-^15^N heteronuclear single quantum coherence (HSQC) NMR experiments were performed. We previously reported the backbone assignments for the BRPF1 bromodomain accomplished by means of multidimensional NMR spectroscopy^31^. Here, the amide-NH chemical shift perturbations induced in the residues of the BRPF1 bromodomain sample were monitored upon the addition of acetylated histone peptides H4K5ac (1-10), H4K8ac (1-10), H4K12ac (4-17), H4K5ac (1-15) as well as the di-acetylated histone ligands H4K5acK8ac (1-10) and H4K5acK12ac (1-15) at increasing concentrations until the saturation point was observed. The bar graphs in **Figures 2A-F** show the normalized chemical shift changes of individual residues in the BRPF1 bromodomain upon addition of the respective mono- and di-acetylated histone peptides, in a 1:5 molar ratio. The affected amino acids were then mapped onto the surface of the apo BRPF1 bromodomain crystal structure (PDB ID: 2D9E), and it was observed that the majority of the altered resonances with large chemical shifts correspond to residues clustered around the acetyllysine binding pocket (**Figure 2A-F**). Large chemical shift perturbations for residue N83 in the BRPF1 bromodomain were consistently observed in our NMR titration studies confirming the importance of this residue in the coordination of acetylated H4 histone peptides. Residues I27, L34, E36, V37, N83, and I88 also showed large chemical shift perturbations upon addition of mono- and/or di-acetylated peptides. A few of the chemical shift perturbations corresponded to residues located more peripherally to the binding pocket. Amino acids including L9, I10, L11, Q55, and N56, located distal to the acetyllysine binding pocket, demonstrated chemical shift perturbations due to conformational changes within the bromodomain that occur in response to ligand binding.

**Figure 2.**
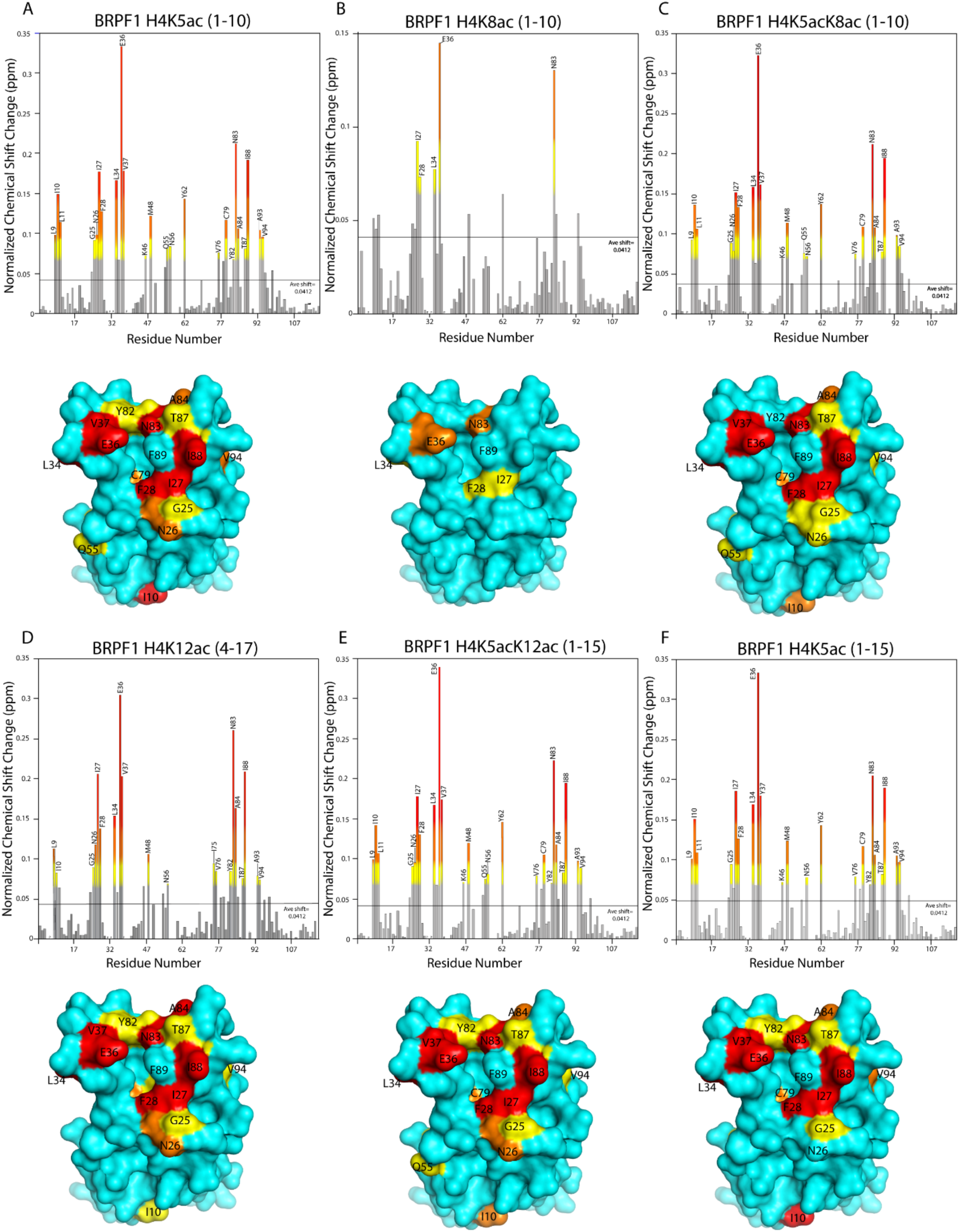
Mapping of the mono- and di-acetylated histone ligand binding interfaces on the BRPF1 bromodomain. (A-F). Histograms show the normalized ^1^H-^15^N chemical shift changes in the backbone amides of the BRPF1 bromodomain in the presence of H4K5ac (1-10) (A), H4K8ac (1-10) (B), H4K5acK8ac (1-10) (C), H4K12ac (4-17) (D), H4K5acK12ac (1-15) (E), and H4K5ac (1-15) (F) in a 1:5 molar ratio. Chemical shift changes 0.5, 1, and 2 standard deviations away from the average shift change are colored yellow, orange, and red, respectively. Amino acid residues with significant chemical shift perturbations upon addition of the histone ligands are mapped onto the surface of the apo structure of the BRPF1 bromodomain (PDBID: 2D9E).

Minimal chemical shift perturbations were seen with the H4K8ac (1-10) histone ligand with only G36 and N83 showing significant perturbations; this is consistent with the weak binding of the H4K8ac ligand to the BRPF1 bromodomain (K_D_ = 168.4 ± 4.7 µM) found by our ITC experiment (**Table 1 and Supplementary Figure S1**). Interestingly, our NMR titration analysis displayed a similar trend in the chemical shift perturbations observed for the mono-versus the di-acetylated histone ligands. The reason for this could be two-fold. Firstly, in the present *in-vitro* condition, the BRPF1 bromodomain shows a similar binding pocket for the mono-versus the di-acetylated H4 histone ligands, thus indicating that there may be other mechanisms involved in more specific interactions of the BRPF1 bromodomain with the di-acetylated H4 histone tail *in-vivo*. BRD7, which shows high structural similarity to BRPF1, has also been reported to lack specificity toward histone ligands under *in-vitro* conditions^40^. Secondly, the second acetyllysine is likely making transient side chain interactions with certain residues surrounding the canonical BRPF1 bromodomain binding pocket not readily detected by our backbone amide NMR titrations. A similar observation was made for the BRDT BD1 bromodomain, which interacts with the di-acetylated H4K5acK8ac histone ligand via the methyl groups of the hydrophobic residues lining the bromodomain pocket^36^.

### Mutational analysis identifies residues that may coordinate the second acetyllysine moiety

To obtain more information about coordination of di-acetyllysine by the BRPF1 bromodomain we used site-directed mutagenesis coupled to ITC binding assays (**Table 2 and Supplementary Figure S2**). To investigate whether both acetyllysine groups in the di-acetylated histone ligands are accommodated in the canonical BRPF1 bromodomain binding pocket, or if secondary binding contacts exist on the BRPF1 bromodomain, we selected six amino acid residues (T24, N26, E30, E36, K81, R91) to mutate to alanine. These experiments were designed to test our hypothesis that if the second acetyllysine group is coordinated via specific side-chain interactions on the BRPF1 bromodomain, then mutating it should decrease the binding affinity to what is seen for the singly acetylated histone ligands. We performed circular dichroism (CD) experiments on the mutant BRPF1 bromodomain proteins (**Table 3 and Figure 3**) to ensure they were properly folded, and we ran ITC binding experiments to test the binding affinities of the BRPF1 bromodomain mutants in combination with H4K5ac (1-10), H4K12ac (1-15), H4K5acK8ac (1-10) and H4K5acK12ac (1-15) histone ligands. The binding affinities with observed stoichiometries of the mutant BRPF1 bromodomains for the histone ligands tested are shown in **Table 2**. The ITC traces of the mutant BRPF1 bromodomains with their histone ligands are shown in **Supplemental Figure S2**. Interestingly, all of the alanine mutations to the BRPF1 bromodomain caused an increase in binding affinity for the H4K5acK12ac ligand, while most of these mutations caused at least a slight decreased in binding affinity for H4K5acK8ac. Our mutational study did not cause the binding affinity of di-acetylated ligands to revert to singly acetylated ligand values, but the T24A, E36A, and K81A mutations did show a weakened binding interaction between the BRPF1 bromodomain and the H4K5acK8ac ligand. The N26A mutation decreased the affinity for the H4K12ac (1-15) ligand, and increased the binding affinity for the H4K5acK12ac (1-15) and H4K5ac (1-10) ligands, but it did not have a large effect on binding of the H4K5acK8ac (1-10) ligand. The remaining mutations either caused no significant difference in the binding interaction or resulted in increased binding affinities with the BRPF1 bromodomain. The inconsistent binding affinity of BRPF1 bromodomain mutants suggests that multiple residues are involved in coordinating the second acetyllysine group.

**Table 2.** Dissociation constants of the wild-type and mutant BRPF1 bromodomain proteins with histone ligands as measured by ITC experiments.

| BRPF1 | H4K5ac (1-10), N | H4K12ac (1-15), N | H4K5K8ac (1-10), N | H4K5K12ac (1-15), N |
| --- | --- | --- | --- | --- |
| Wild type | 60.1 ± 6.8, 0.9 ± 0.1 | 97.3 ± 4.0, 1.0 ± 0.1 | 26.7 ± 1.8, 1.0 ± 0.0 | 52.0 ± 2.9, 1.0 ± 0.0 |
| T24A | 28.9 ± 1.0, 1.0 ± 0.0 | 105.7 ± 8.3, 0.9 ± 0.1 | 99.4 ± 5.3, 1.0 ± 0.1 | 31.2 ± 4.6, 1.0 ± 0.0 |
| E30A | 79.4 ± 5.7, 1.1 ± 0.1 | 104.5 ± 4.0, 1.1 ± 0.0 | 27.2 ± 0.7, 1.0 ± 0.0 | 39.0 ± 1.6, 1 ± 0.0 |
| E36A | 51.4 ± 1.9, 1.0 ± 0.0 | 54.7 ± 5.5, 1.0 ± 0.0 | 43.1 ± 1.4, 1 ± 0.0 | 31.0 ± 2.6, 1.0 ± 0.0 |
| K81A | 59.9 ± 3.8, 1.0 ± 0.0 | 45.0 ± 3.0, 1.0 ± 0.0 | 45.7 ± 5.6, 1 ± 0.0 | 24.6 ± 0.8, 1.0 ± 0.0 |
| R91A | 37.3 ± 1.3, 1.0 ± 0.0 | 79.7 ± 5.1, 1.0 ± 0.0 | 21.5 ± 1.9, 0.9 ± 0.0 | 23.0 ± 3.8, 1.0 ± 0.0 |
| N26A | 44.1 ± 0.9, 1.0 ± 0.0 | 122.3 ± 2.9, 1.2 ± 0.1 | 34.5 ± 1.3, 1.0 ± 0.0 | 34.2 ± 6.8, 1.0 ± 0.0 |

**Table 3:**
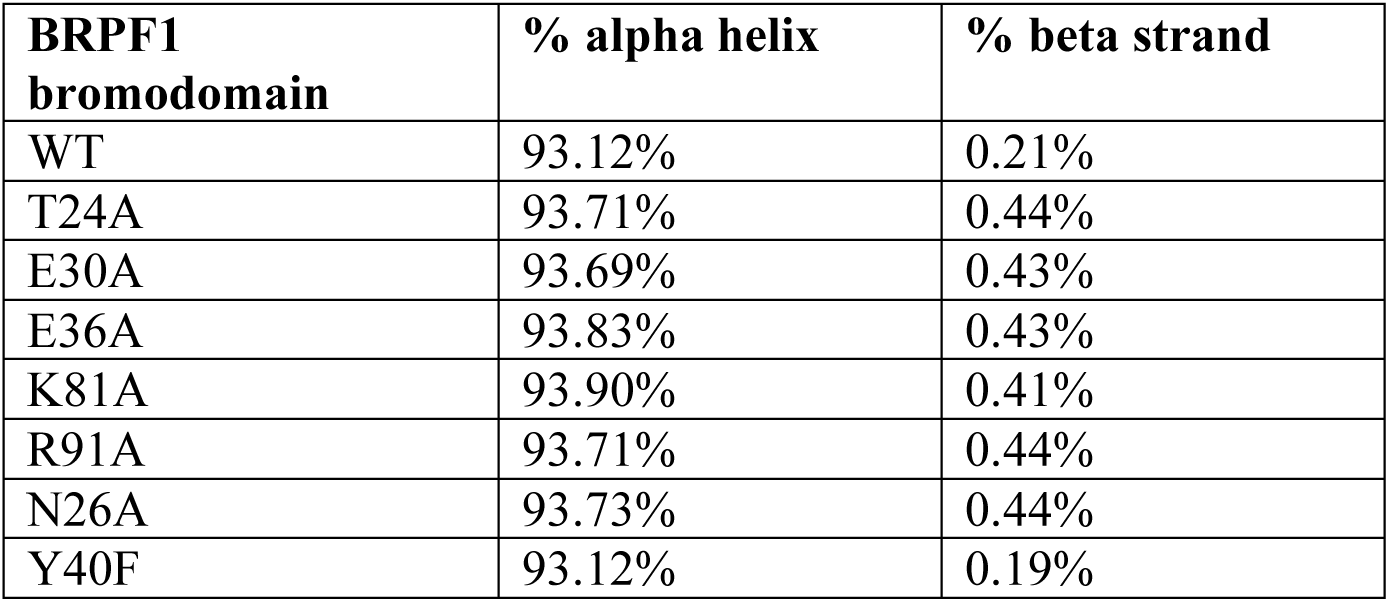
Percentages of α-helix and β-strand composition in wild-type (WT) and mutant BRPF1 bromodomain proteins calculated from circular dichroism experiments.

| BRPF1<br>bromodomain | % alpha helix | % beta strand |
| --- | --- | --- |
| WT | 93.12% | 0.21% |
| T24A | 93.71% | 0.44% |
| E30A | 93.69% | 0.43% |
| E36A | 93.83% | 0.43% |
| K81A | 93.90% | 0.41% |
| R91A | 93.71% | 0.44% |
| N26A | 93.73% | 0.44% |
| Y40F | 93.12% | 0.19% |

**Figure 3.**
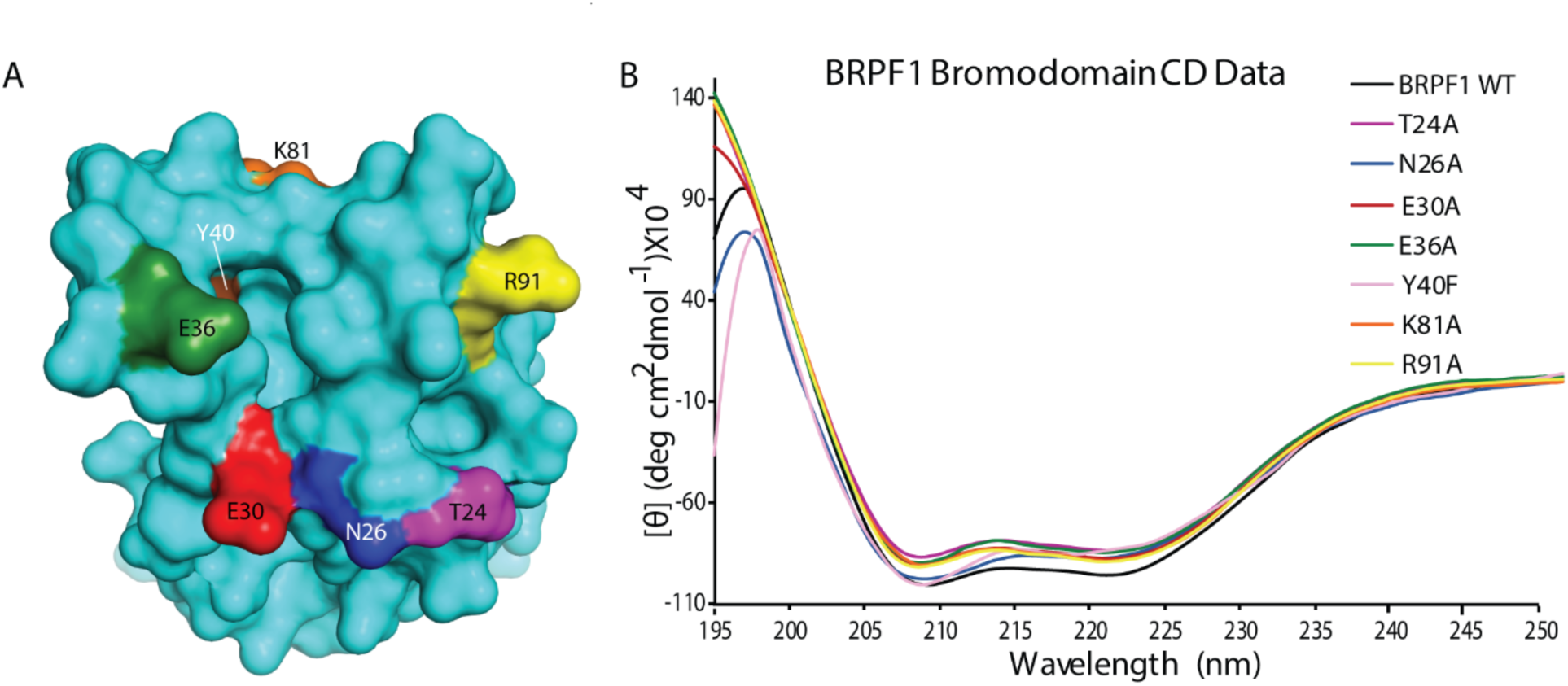
Circular dichroism spectra of the wild-type and mutant BRPF1 bromodomains. (A). Surface representation of the apo structure (cyan) of the BRPF1 bromodomain showing mutated residues. The amino acid residues N26, T24, E30, E36, Y40, K81. and R91 mutated to alanine are indicated as shown. (B). Secondary structure of the BRPF1 bromodomain wild-type and mutant bromodomain proteins measured by circular dichroism. Circular dichroism molar ellipticity plots of the BRPF1 wild type and mutant bromodomains are shown. Each protein is represented by a different color and all show clear α-helical structure to match the wild type BRPF1 bromodomain.

### The role of ordered water in bromodomain structure and histone ligand coordination

We made a conservative change to Y40 by mutating the residue to phenylalanine in order to investigate how loss of water coordination would impact acetyllysine recognition. The BRPF1 Y40F bromodomain variant failed to bind mono- or di-acetylated histone ligands (**Table 2 and Figure 4)**. Thus, the ordered water in the bromodomain binding pocket is important for the recognition of acetylated histone ligands.

**Figure 4.**
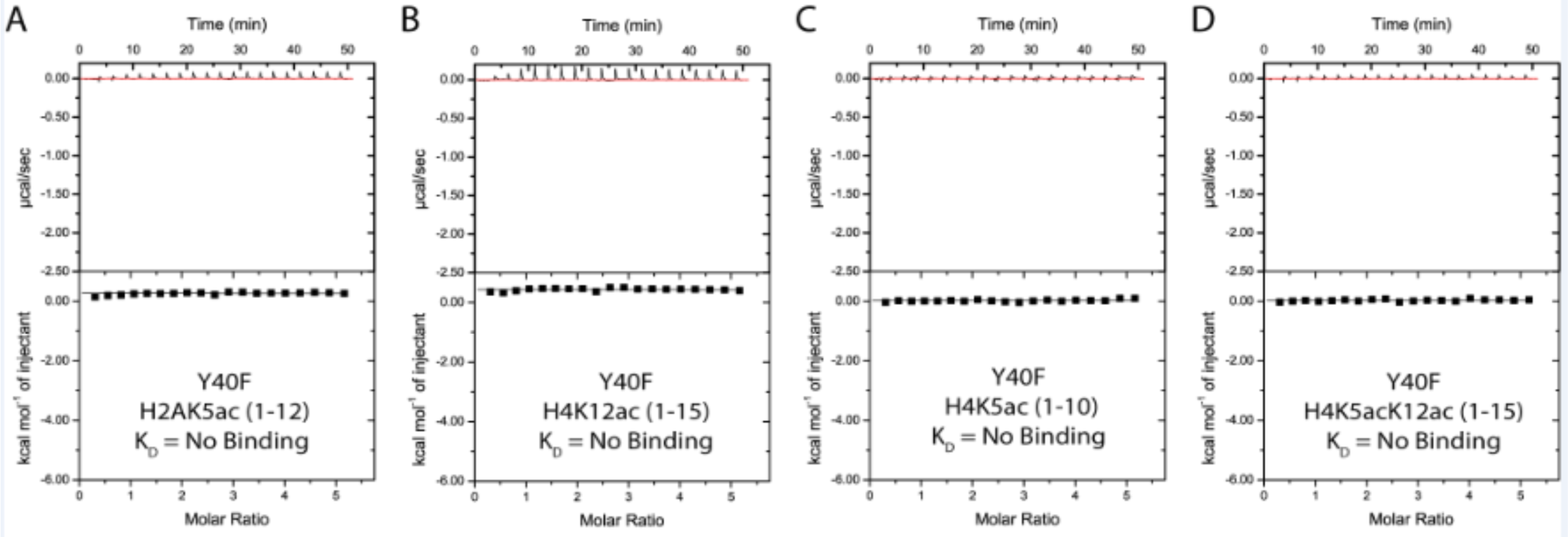
ITC measurements of the Y40F mutant bromodomain involved in water coordination, with acetylated histone peptides. No binding was detected for all the four histone peptides tested with the Y40F mutant bromodomain which shows its important role in water coordination in the bromodomain binding pocket.

## DISCUSSION

Previous studies on the BRPF1 bromodomain revealed that it recognizes several acetylated lysine residues on the N-terminus of histone tails^31^. The BRPF1 bromodomain predominantly selects for acetylation modifications on the histone H4 tail; however, it is also able to recognize the histone H3 and H2A ligands H3K14ac and H2AK5ac^31^. More recently it has been discovered that bromodomain-containing proteins such as BRD2, BRD3, and BRD4 can bind to histone ligands with di-acetyllysine marks^41^. Another example of this is the first bromodomain (BD1) in the bromodomain testis specific protein (BRDT), which is a member of the bromodomain and extra-terminal domain (BET) subfamily. BRDT BD1 failed to recognize mono-acetylated histone H4 tails, but was able to bind multiple di-acetylated histone H4 peptides as well as tetra-acetylated histone H3 peptides^36^. BRDT BRD1 recognizes the di-acetylated histone ligand H4K5acK8ac with a low micromolar affinity, and the crystal structure of this complex demonstrated that the ‘wide pocket’ of the BRDT BD1 bromodomain is able to accommodate both acetylated lysine residues within the canonical bromodomain binding pocket^36^. By contrast the ‘keyhole pocket’ of the yeast GCN5 bromodomain is able to accommodate only one acetyllysine group inside the binding pocket at a time. Coordination of di-acetyllysine by the first bromodomain in the BET subfamily appears to be conserved as demonstrated by ligand binding assays with BRD1 and BRD2^42,43^, as well as by structural models of BRD4 BD1 with H4K5acK8ac and H4K12acK16ac (PDBIDs 3UVW and 3UVX, respectively)^4^. It has been suggested that other bromodomain protein families can recognize di-acetylated histone ligands as well, possibly through alternative binding modes. The structures of the BAZ2A and BAZ2B bromodomains complexed, respectively, with the di-acetylated histone H4 ligands H4K16acK20ac and H4K8acK12ac have been determined (PDBIDs 4QBM and 4QC3)^38^. In these structures, BAZ2A can coordinate the di-acetylated ligand as a monomer, much like the BET family bromodomains, while the BAZ2B bromodomain forms a dimeric complex to coordinate the H4K8acK12ac histone ligand.

In this study, we show that the BRPF1 bromodomain can selectively recognize multiple histone mark combinations and that it binds preferentially to di-acetylated ligands on the N-terminus of histone H4. The results from our ITC binding assays demonstrate that this bromodomain has the highest affinity for the histone H4K5acK8ac ligand, followed by the H4K5acK12ac ligand. Histone H4 acetylation occurs progressively from mono-acetylated H4K16ac, followed by acetylation at the K12, K8, and K5 sites^44,45^. Therefore the presence of K5 and K8 acetylation marks are characteristic of chromatin hyperacetylation *in-vivo*^*46*^. Previous studies also reported that newly synthesized histone H4 has transient acetylation marks on the K5 and K12 positions^37,47^. Given the fact that BRPF1 recruits MOZ to particular sites on active chromatin and remains at chromosomes during mitosis^9^, it is reasonable to assume that the BRPF1 bromodomain plays an important role in the recognition of hyperacetylation PTMs on chromatins during cell division. BRPF1 binding would further drive acetylation of the histones via the acetyl transferase activity of MOZ, which is activated by BRPF1^13^. Since BRPF1 has been shown to play an important role in cell proliferation^11^, it is possible that the recognition of these di-acetylation marks, characteristic of dividing cells, is driving cancer progression.

We also tested the histone H4K5acK16ac and H4K8acK16ac ligands and found that these di-acetyllysine combinations weakened the BRPF1 bromodomain interaction with the histone tail. Examination of the histone H3/H4 proteoforms found in human cancer cell lines revealed that in promyelocytic leukemia cells (HL60) the H4K5acK16ac mark is down-regulated, while it is up-regulated in acute promyelocytic (NB4) cells^48^. Additionally, the H4K8acK16ac mark was down-regulated in all leukemia cell lines tested. Thus, as histone PTMs are altered in disease states, the availability of particular acetyllysine combinations will enhance or inhibit the interaction of BRPF1 with chromatin and influence recruitment of the MOZ/MORF HAT complex.

Results from our previous study demonstrated that the BRPF1 bromodomain recognizes the singly-acetylated histone ligands H2AK5ac, H4K12ac, H4K8ac, H4K5ac, and H3K14ac respectively^31^. Our current results demonstrate that of the singly acetylated histones, the BRPF1 bromodomain binds preferentially to H4K5ac, which is likely the predominant interaction driving recognition of di-acetylated ligands. Interestingly, the ITC binding data show that the BRPF1 bromodomain functions as a di-acetyllysine reader, with a binding pocket able to recognize PTMs over span of 8-10 residues on each histone tail. Furthermore, our results show that the interaction of the BRPF1 bromodomain with histone ligands is enthalpy driven. A similar result was also reported for the BRPF1 bromodomain with small-molecule ligands, and with ATAD2 bromodomain-ligand complexes^49,50^. Additionally, the BRPF1 bromodomain interacts with mono- and di-acetylated histone ligands as a monomer in solution based on the binding stoichiometry (N: peptide-to-bromodomain ratio) observed in the ITC binding assays and confirmed in AUC experiments.

This stoichiometry differs from that observed for the family IV BRD9 bromodomain, which binds to the histone H4K5acK8ac ligand with an ITC N-value of 0.44, suggesting a dimeric interaction with two bromodomains bound to one histone peptide^19^. The family IV bromodomains possess a “keyhole style” pocket that, owing to the presence of a large aromatic gatekeeper residue, can only accommodate one acetyllysine group in the canonical binding site^19,36^. Thus, in ITC binding assays and crystallization studies the BRD9 bromodomain interacts with the di-acetylated histone ligand by coordinating the H4K5ac group in one bromodomain with the H4K8ac site bound to a second bromodomain module (PDBID: 4YYI)^19^. A related structure shows that the BRPF1 bromodomain binds to the histone H4K5acK8ac in an identical manner as an apparent dimer (PDBID: 5FFW)^12^. However, this is likely due to crystal packing effects caused by the crystallization conditions used for solving the crystal structure, since our ITC binding data and analytical ultracentrifugation analysis of the BRPF1 bromodomain support monomeric coordination of both mono- and di-acetylated histone ligands in solution. Also, unlike the BET proteins, which contain two adjacent bromodomain modules^51^, the BRPF1 protein has only one bromodomain, and this protein functions as a single subunit within the MOZ/MORF HAT complexes^4,10^. Thus, the BRPF1 bromodomain is likely to be monomeric in its biologically active form.

Histone H4 can be post-translationally modified at multiple positions and is frequently acetylated at lysine 5, 8, 12, and 16. It also often carries a phosphorylation mark on serine 1, as well as methyl groups on arginine 3 and lysine 20^52^. The SET-domain methyltransferases performs methylation of lysine on the histone tails^53^, which can be mono-, di- or tri-methylated. Lysine at position 5 and 12 of histone H4 can also be methylated^32,54^. Since acetylation at lysine 5 and 12 on histone H4 are the positions the BRPF1 bromodomain binds with highest affinity, we tested the effect tri-methylation at these sites would have on bromodomain binding. Our results demonstrated that tri-methylation of H4K5 reduced the binding affinity of the BRPF1 bromodomain for the H4K12ac ligand, while tri-methylation of H4K12 did not have a significant impact on the recognition of H4K5ac. We also characterized how phosphorylation of serine 1 modulates the acetyllysine binding activity of the BRPF1 bromodomain and showed that this mark has an inhibitory effect on recognition of H4K5ac. Histone phosphorylation has previously been linked to DNA damage repair mechanisms^55^, and H4S1phos is known to be enriched 20-to 25-fold in nucleosomes that are located adjacent to double strand breaks^56^. The impact of modifications adjacent to acetyllysine recognition sites may be biologically relevant to BRPF1’s function in recruiting the MOZ HAT tetrameric complex to distinct sites on chromatin depending on the epigenetic landscape. For example, phosphorylation of histone H4S1 at sites of DNA damage could function to prevent or delay the transcriptional activation signaling by acetylation modifications until the genetic information can be repaired. Therefore, our study represents an important first step in understanding how the BRPF1 bromodomain is able to recognize and bind di-acetyllysine on the histone tail, and how cross-talk among adjacent marks influences its activity.

To delineate the acetyllysine binding site on the BRPF1 bromodomain, we used NMR chemical shift perturbation data to identify the residues involved in di-acetyllysine recognition, and to observe whether residues outside the canonical acetyllysine binding pocket are important for coordinating the second acetyllysine modification. We introduced additional mutations into the BRPF1 bromodomain binding pocket to test hypotheses based on previous studies of acetyllysine coordination by other related bromodomains^57-60^. I27 is found in the BRPF1 “NIF Shelf”, which is important for ligand selectivity^61^. L34, E36, and V37 are all located in the flexible ZA loop adjacent to the acetyllysine binding site, which is predicted to be important for increasing the on-rate of histone tail binding^60^. N83 is the universally conserved asparagine residue involved in coordination of the acetyllysine modification on histones^61^. I88 is adjacent to the ‘gatekeeper’ residue F89 (unassigned) that lines the acetyllysine binding pocket. Residue E36 in the BRPF1 bromodomain showed the largest chemical shift perturbation upon addition of either mono- or di-acetylated histones. Alignment of the apo crystal structure of the BRPF1 bromodomain (PDBID: 2D9E) with BRPF1 bromodomain structures in combination with the H2AK5ac and H4K12ac ligands (PDBIDs: 4QYL and 4QYD)^59^, demonstrate that the E36 residue does not contact the histone ligands directly, but undergoes a large conformational change as part of the ZA loop to accommodate peptide binding. Mutation of E36 to alanine did not significantly alter the binding affinity of the BRPF1 bromodomain with the H4K5ac (1-10) ligand, but it decreased the interaction of the BRPF1 bromodomain with the di-acetylated histone H4K5acK8ac (1-10) ligand (**Table 2**), suggesting it may be an important contact for the K8ac moiety. A recent study investigating the intermolecular contacts between the related ATAD2 family IV bromodomain and the H4K5acK12ac ligand found that residue E1017, which is equivalent to E36 in the BRPF1 bromodomain, comes within 5 Å of H4K8 in molecular dynamic simulations of the holo complex^60^.

A prior study on the ATAD2 bromodomain found the H3K14ac ligand bound to a secondary site distal from the canonical bromodomain binding site^57^. In this ATAD2 structure (PDBID: 4TT4), two residues were involved in coordination of the acetyllysine moiety, R1005 and H1076. H1076 is equivalent to T24 in the BRPF1 bromodomain. The T24A mutant showed a two-fold stronger binding affinity for the H4K5ac (1-10) ligand than the wild-type BRPF1 bromodomain, while also having a weaker interaction with the H4K5acK8ac (1-10) ligand. This suggests that residue T24 plays some role in coordinating either the K5ac or the K8ac in the di-acetylated histone ligand. Notably, mutation of either K81 to alanine significantly weakened the interaction of the BRPF1 bromodomain for the H4K5acK8ac ligand, also indicating a role for this residue in coordination of this di-acetylated histone ligand. However, the N26A and E30A mutations resulted in modest changes in the binding affinity of the BRPF1 bromodomain for the histone ligands, suggestive of the neutral role of these residues in acetyllysine binding. Similarly, the R91A mutation resulted in slightly stronger binding interactions of the BRPF1 bromodomain with both the mono- and di-acetyllysine ligands. When the histone peptides contain two acetyllysine groups, it is possible for either group to be inserted into the canonical bromodomain binding pocket. One potential reason we observed an increase in binding affinity of the mutant BRPF1 bromodomain proteins with the di-acetyllysine ligands could be because their binding is now stabilized in one orientation over the other. In a di-acetylated ligand, not only is each acetyllysine group able to insert into the canonical binding pocket, but the peptide can bind in different orientations (e.g. from N-to C-terminal or vice versa) across the surface of the bromodomain binding pocket. Thus, by altering the BRPF1 bromodomain surface contacts with histone residues surrounding the primary acetyllysine inserted into the canonical binding pocket, some of our mutations may be preferentially selecting for one orientation over another, thereby stabilizing the ligand binding interaction with both mono- and di-acetylated histone peptides. Overall, our NMR titration and mutagenesis data demonstrated that the acetyllysine binding pocket in BRPF1 bromodomain binds a variety of acetylated histone ligands with a conserved binding mechanism, and that increased sidechain and/or hydrophobic interactions are involved in the coordination of the second acetyllysine group, thus resulting in stronger binding affinities for the di-acetylated histone ligands.

Previous studies have identified ordered waters in the bromodomain binding pocket that coordinate the acetyllysine modification^59,62^. One of these waters (w1) connects the carbonyl oxygen of the acetyllysine moiety to the hydroxyl group of Y40. Y40 is located in the αAZ loop of the bromodomain module and by sequence alignment is conserved in all but two of the human bromodomain proteins^4^. Thus, this residue is even more highly conserved than the binding site asparagine residue that contacts the acetyllysine group directly. Our conservative mutation of this residue to phenylalanine demonstrated that Y40 is crucial for water coordination and sheds light on why Y40 is the most highly conserved amino acid residue found in human bromodomains.

In summary, our results demonstrated that the BRPF1 bromodomain can recognize multiple acetyllysine combinations on the histone tail. The BRPF1 bromodomain preferentially selects for the di-acetyllysine modifications H4K5acK8ac and H4K5acK12ac, which are associated with hyperacetylation and cell division, respectively^37,46,47^. Our results also confirm that cross-talk between PTMs on histone tail peptides can modulate acetyllysine recognition by the BRPF1 bromodomain. This research provides further insights into how the BRPF1 bromodomain recruits its partners in the MOZ/MORF HAT complexes to chromatin, with different combinations of PTMs either promoting or inhibiting the binding interaction. Chromosomal translocations of the MOZ HAT are associated with acute myeloid leukemia, and fusion of MOZ with TIF2, CBP and p300 are thought to drive leukemogenesis through aberrant acetylation^63-65^. Understanding how cross-talk influences bromodomain binding specificity and affinity is an important first step to connect how epigenetic changes contribute to the etiology of disease and may provide new mechanisms for therapeutic intervention in the treatment of AML and other diseases.

## MATERIALS AND METHODS

### Plasmid construction

The cDNA coding for human BRPF1 was kindly provided to us by Dr. Xiang-Jiao Yang at McGill University. Residues 629-742 encoding the BRPF1 bromodomain region were amplified using PCR and subsequently cloned into the pDEST15 vector encoding an N-terminal glutathione S-transferase (GST) tag. The Gateway cloning technology (Invitrogen) was used for cloning the BRPF1 bromodomain as described previously^31^. BRPF1 bromodomain mutants T24A, N26A, E30A, E36A, Y40F, K81A, and R91A were generated using the QuikChange^®^ mutagenesis procedure (Stratagene) as described previously^66^. All of the mutants were also cloned into the pDEST15 vector and the DNA sequences of the wild-type (WT) and mutant proteins were verified by Eurofins Genomics. The WT and mutant plasmids were then transformed into *Escherichia coli* Rosetta™ 2(DE3)pLysS competent cells (Novagen) for protein expression.

### BRPF1 bromodomain expression and purification

*E. coli* Rosetta™ 2(DE3)pLysS cells expressing the WT and mutant GST-tagged BRPF1 bromodomain were grown in terrific broth (TB) at 37°C until an OD_600_ of 1.0 was reached. The temperature was reduced to 16°C for 1 h, and the cultures were induced by 0.25 mM isopropyl β-D-1-thiogalactopyranoside (IPTG) and incubated overnight for over 16 h at 16°C. The cells were harvested by centrifugation and sonicated after resuspension in lysis buffer (50 mM Tris pH 7.5, 150 mM NaCl, 0.05% Nonidet P-40, and 1mM DTT) containing 0.1 mg/mL lysozyme (Thermo Scientific). After centrifugation at 10,000 RPM, the cell supernatant was mixed with glutathione agarose resin (Thermo Scientific) and incubated on ice (4°C) for 2 h while agitating. The BRPF1 bromodomain protein was purified on glutathione agarose resin in a 25 mL Econo-Column^®^ (Bio-Rad) by first washing the beads extensively with wash buffer containing 20 mM Tris pH 8.0, 150 mM NaCl, and 1 mM DTT. The GST tag was cleaved overnight at 4°C by addition of PreScission Protease (GE Healthcare), and the BRPF1 bromodomain was eluted with wash buffer. The eluted fractions were pooled, concentrated to approximately 3 mL, and dialyzed into appropriate buffers. For NMR experiments the ^15^N-labeled BRPF1 bromodomain was expressed and purified as previously described^31^. The uniformly ^15^N-labeled BRPF1 bromodomain was prepared at a concentration of 0.4 mM in 20 mM Tris-HCl pH 6.8, 150 mM NaCl, and 10 mM DTT with 10% D_2_O added to the sample. For ITC and circular dichroism (CD) experiments, the purified BRPF1 bromodomain was dialyzed, respectively, into ITC buffer (20 mM NaH_2_PO_4_ pH 7.0, 150 mM NaCl, and 1 mM TCEP), or CD buffer (50 mM NaH_2_PO_4_ pH 7.0 and 50 mM NaCl). For analytical ultracentrifugation (AUC) experiments, the protein was further purified over a gel filtration column (HiPrep 16/60 Sephacryl S-100 High Resolution, GE Healthcare) equilibrated with AUC buffer (50 mM NaH_2_PO_4_ pH 7.0, 50 mM NaCl, and 1 mM TCEP) via fast-protein liquid chromatography. The concentration of the BRPF1 bromodomain was calculated from optical absorbance measurements with the A_280_ = 7450 M^−1^cm^−1^. The purity of the BRPF1 bromodomain was verified by SDS-polyacrylamide gel electrophoresis stained with GelCode Blue Safe protein stain (Thermo Scientific).

### Histone peptide synthesis

Histone peptides containing unique acetylated and/or methylated lysine, as well as phosphorylated serine modifications were synthesized by GenScript and purified by HPLC to 98% purity. The correct composition of each peptide was confirmed by mass spectrometry.

### Isothermal titration calorimetry

ITC measurements were conducted using a MicroCal iTC200 instrument (Malvern Panalytical). All experiments were performed at 5°C. The BRPF1 WT and mutant bromodomain proteins were dialyzed into ITC buffer containing 20 mM NaH_2_PO_4_ pH 7.0, 150 mM NaCl, and 1 mM TCEP for 48 hours before experiments were performed. All titration experiments were performed as follows: 200 µM of the BRPF1 bromodomain protein was injected into the sample cell using a loading syringe, with 5 mM of histone peptide loaded into the injection syringe. The sample cell received one preliminary injection of 0.5 µL of the histone peptide followed by 19 injections of 2 µL histone peptide. For all titration experiments, each injection was spaced at 150 s time intervals after an initial delay of 60 s, and the injection syringe was stirred continuously at 1000 RPM. The preliminary injection was excluded prior to integration of the binding curve and calculation of the dissociation constants (K_D_s). First, the raw data were corrected for non-specific heats of dilution determined by the magnitude of the peaks appearing after the system reaches complete saturation and normalized for concentration. The raw data were then integrated using a Marquandt nonlinear least-squares analysis according to a 1:1 binding model, normalized for concentration, and analyzed assuming a single set of identical binding sites to calculate the binding affinity, stoichiometry (N), and the thermodynamic values. Binding data for all experiments were analyzed using the ORIGIN 7.0 software (OriginLab Corporation). All experiments where binding occurred were performed in triplicate (with the exception of H3K14ac (1-24), which was performed in duplicate), while nonbinding experiments were performed in duplicate and standard errors were reported as standard deviations.

### Circular dichroism spectroscopy

Circular Dichroism (CD) spectra were recorded on a JASCO J-815 CD Spectrometer (JASCO, Japan) at 25°C in a 1.6 cm cell. BRPF1 bromodomain WT and mutant proteins were dialyzed in CD buffer containing 50 mM NaH_2_PO_4_ pH 7.0 and 50 mM NaCl and then diluted to between 0.1 µM and 0.5 µM concentration. CD spectra were measured from 199 nm to 260 nm. Two spectra were measured and averaged for each WT and mutant bromodomain protein sample. The spectra were analyzed using the K2D3 structure prediction software^67,68^ to determine the percent α-helical and β-sheet content for the WT and each of the mutant BRPF1 bromodomain proteins.

### NMR spectroscopy

Chemical shift perturbation experiments were performed using 0.4 mM of uniformly ^15^N-labeled BRPF1 bromodomain in NMR buffer containing 20 mM Tris-HCl pH 6.8, 150 mM NaCl, 10 mM DTT and 10% D_2_O. Titration mixtures of the BRPF1 bromodomain protein and each of the modified histone peptides (H4K5acK8ac residues 1-10, H4K5acK12ac (1-15), H4K5ac (1-10) or 1-15 and H4K8ac (1-10)) were prepared at concentration ratios of 1:0, 1:0.6, 1:1.2, 1:3 and 1:6 in a volume of 150 µL. The sample mixtures were then transferred into 3-mm NMR tubes (Bruker). Two-dimensional ^15^N HSQC (heteronuclear single quantum coherence) experiments for all samples were run at 25°C and NMR data collected as described previously^31^. Normalized chemical shift changes were calculated as described in^69^ using the equation:

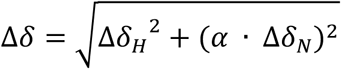

where *α* = 0.14 for all residues except glycine, which has *α* = 0.2, and Δδ_H_ and Δδ_N_, respectively, are the changes in proton and nitrogen chemical shift in ppm. To map the chemical shift perturbations onto the BRPF1 bromodomain surface, we calculated the standard deviation (σ) of the Euclidean chemical shift change over all residues of all bromodomain:peptide combinations. Residues whose chemical shift change was greater than 0.5σ were colored yellow, while residues with chemical shift changes greater than 1σ and 2σ were colored orange and red, respectively. The backbone resonance assignments for the BRPF1 bromodomain were completed as described previously^31^.

### Analytical ultracentrifugation

For all AUC experiments, samples of the WT and mutant BRPF1 bromodomain proteins were prepared at an OD_280_ of 0.6 (80.5 μM) in AUC buffer containing 50 mM NaH_2_PO_4_ pH 7.0, 50 mM NaCl, and 1 mM TCEP. The BRPF1 bromodomain protein alone, and in combination with unmodified histone H4 peptide residues 1-15 mixed in a 1:1 molar ratio, were included as controls. All other acetylated histone peptides (H4K5ac 1-15, H4K12ac 1-15, H4K8ac 1-10, H4K5acK8ac 1-10, and H4K5acK12ac 1-15) tested with the BRPF1 bromodomain were also mixed in a 1:1 ratio. AUC measurements were performed at 20°C by UV intensity detection in a Beckman Optima AUC at the Center for Analytical Ultracentrifugation of Macromolecular Assemblies at the University of Texas Health Science Center at San Antonio and at the Canadian Center for Hydrodynamics at the University of Lethbridge, Alberta, Canada, using an An50Ti rotor and standard 2-channel epon centerpieces (Beckman-Coulter). All the samples were measured in the same run and at multiple wavelengths (278 nm, 280 nm, and 282 nm), and the concentrations measured were reported in 280 nm absorbance units. All the AUC data were analyzed with UltraScan-III ver. 4.0, release 2480^70^. Hydrodynamic corrections for buffer density and viscosity were estimated by UltraScan to be 1.006 g/mL and 1.0239 cP. The partial specific volume of the BRPF1 bromodomain was estimated by UltraScan from protein sequence analogous to methods outlined in Laue et al. Sedimentation Velocity (SV) data were analyzed according a published method^71^. Optimization was performed by two-dimensional spectrum analysis (2DSA)^72^ with simultaneous removal of time- and radially-invariant noise contributions^73^. After inspection of the 2DSA solutions, the fits were further refined using the parametrically constrained spectrum analysis (PCSA)^74^, coupled with Monte Carlo analysis to measure confidence limits for the determined parameters^75^. The calculations are computationally intensive and were carried out on high performance computing platforms^76^. All calculations were performed on the Lonestar cluster at the Texas Advanced Computing Center (TACC) at the University of Texas at Austin or on XSEDE clusters at TACC or at the San Diego Supercomputing Center.

## AUTHOR INFORMATION

### Author Contributions

**The manuscript was written through contributions of all authors. All authors have given approval to the final version of the manuscript.**

## ACKNOWLEDGMENTS

We are thankful to Dr. X.J. Yang at McGill University for providing us with the isolated cDNA from human BRPF1 (Gene ID:7862).

## ABBREVIATIONS

AML: acute myeloid leukemia
AUC: analytical ultracentrifugation
BRPF1: bromodomain-PHD finger protein 1
CD: circular dichroism
DTT: dithiothreitol
HAT: histone acetyltransferase
MEAF6: MYST/Esa1-associated factor 6
ING5: inhibitor of growth 5
ITC: isothermal titration calorimetry
MOZ: monocytic leukemic zinc-finger
NMR: nuclear magnetic resonance
PCR: polymerase chain reaction
PHD: plant homeodomain
PTMs: post-translational modifications
PWWP: proline-tryptophan-tryptophan-proline
SV: sedimentation velocity
TCEP: tris(2-carboxyethyl)phosphine).

