## Supplemental Figures for "The BRPF1 bromodomain is a molecular reader of di-acetyllysine"

### SUPPLEMENTARY FIGURES

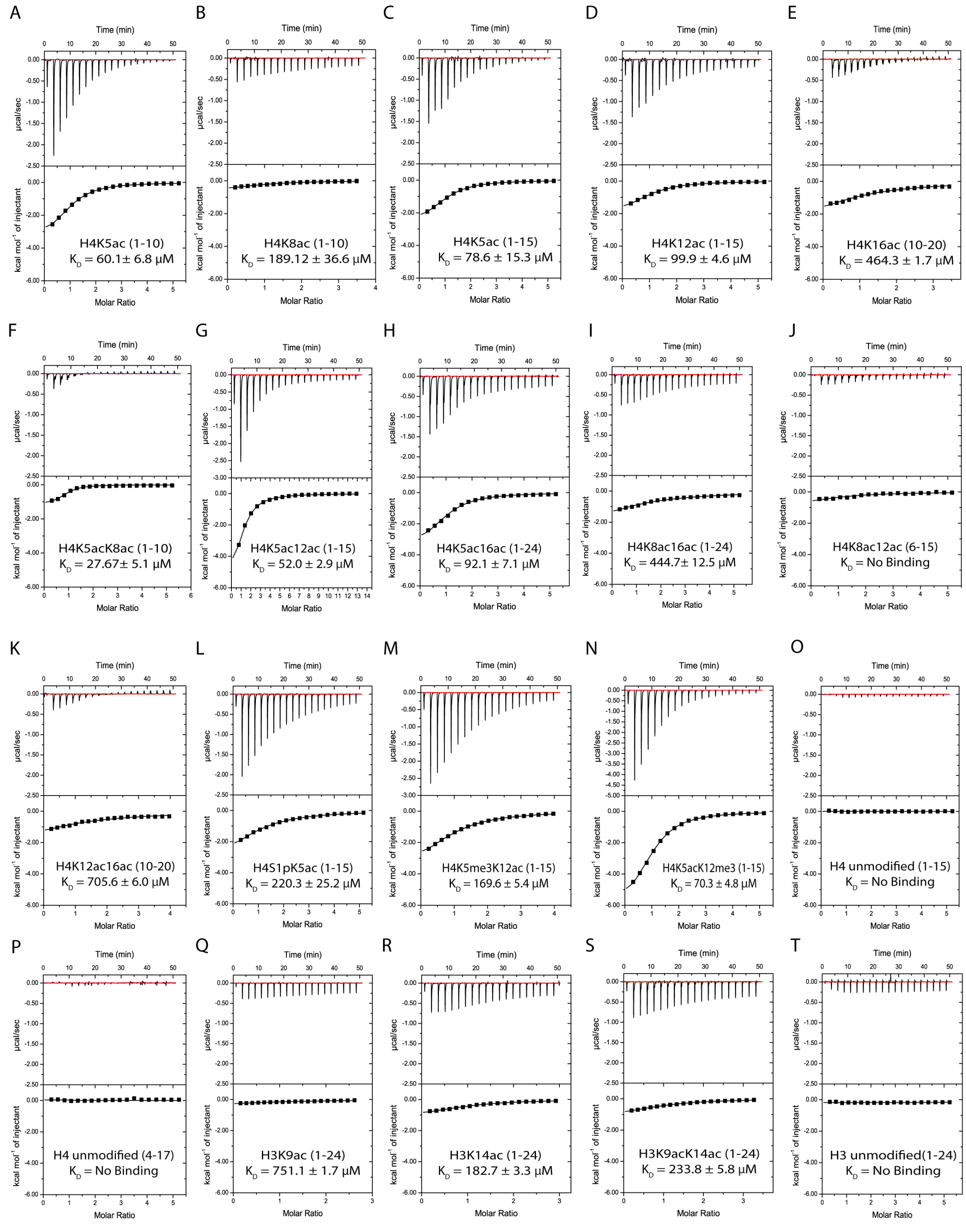


**Supplemental Figure S1: ITC measurements of the interactions between the BRPF1 bromodomain and acetylated histone ligands.** (A-P). Exothermic ITC enthalpy plots for the binding of the BRPF1 bromodomain to mono-, di- and unacetylated histone H4 peptides. The calculated dissociation constants are indicated for each peptide tested.

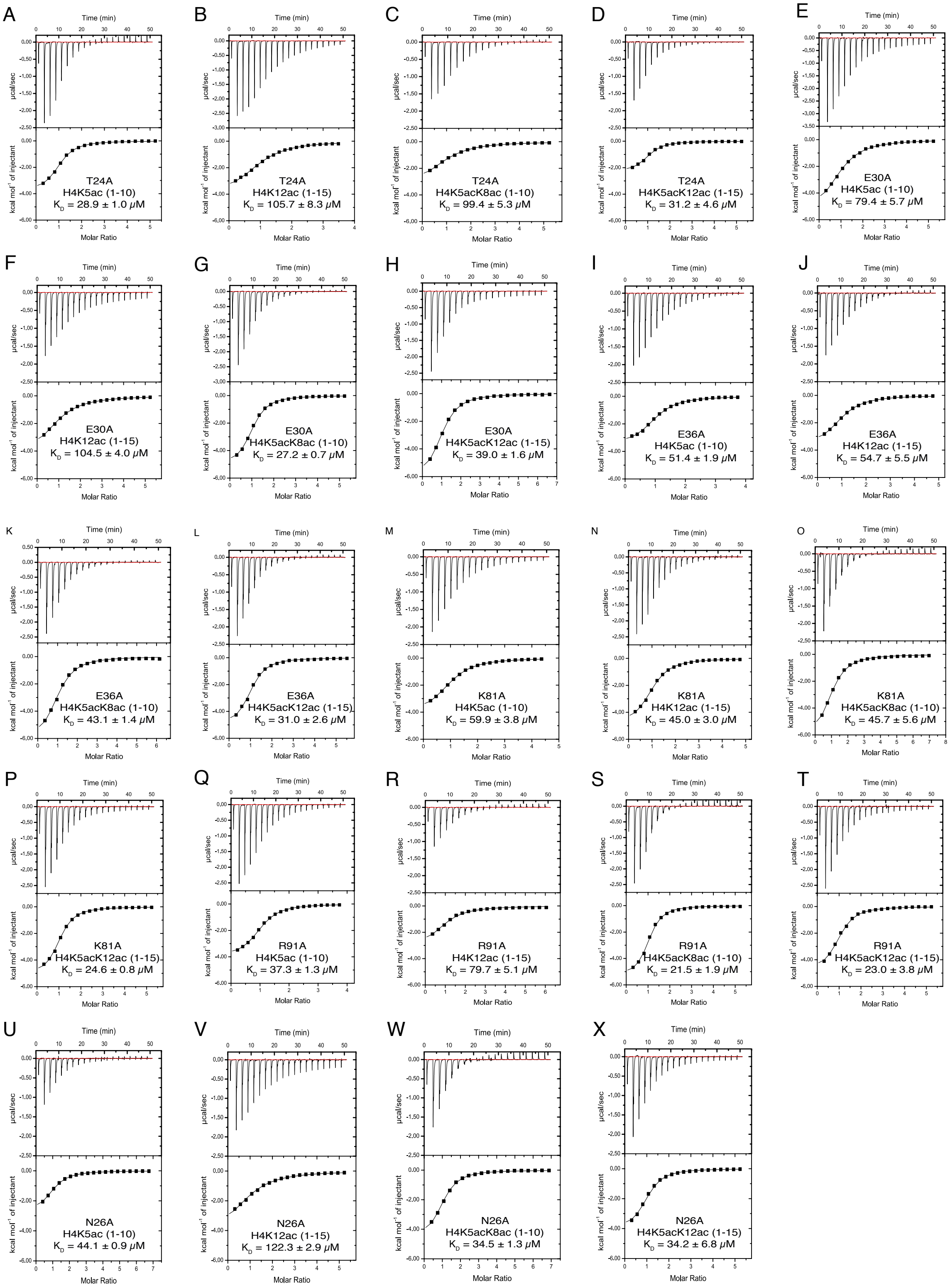


**Supplemental Figure S2**: **ITC measurements of the interactions between the BRPF1 bromodomain mutant proteins and acetylated histone ligands.** (A-X). Exothermic ITC enthalpy plots for the binding of mutant BRPF1 bromodomain proteins to mono- and di-acetylated histone H4 peptides. The calculated dissociation constants are indicated for each peptide tested.
